# SEMplMe: A tool for integrating DNA methylation effects in transcription factor binding affinity predictions

**DOI:** 10.1101/2020.08.13.250118

**Authors:** Sierra S. Nishizaki, Alan P. Boyle

## Abstract

**Motivation:** Aberrant DNA methylation in transcription factor binding sites has been shown to lead to anomalous gene regulation that is strongly associated with human disease. However, the majority of methylation-sensitive positions within transcription factor binding sites remain unknown. Here we introduce SEMplMe, a computational tool to generate predictions of the effect of methylation on transcription factor binding strength in every position within a transcription factor’s motif.

**Results:** SEMplMe uses ChIP-seq and whole genome bisulfite sequencing to predict effects of methylation within binding sites. SEMplMe validates known methylation sensitive and insensitive positions within a binding motif, identifies cell type specific transcription factor binding driven by methylation, and outperforms SELEX-based predictions for CTCF. These predictions can be used to identify aberrant sites of DNA methylation contributing to human disease.

**Availability and Implementation:** SEMplMe is available from https://github.com/Boyle-Lab/SEMplMe.

**Supplementary Information:** Supplementary data are available at *Bioinformatics* online.

## Introduction

DNA methylation is an epigenetic mark as it contributes to changes in the information content of DNA without changing the underlying sequence. The majority of DNA methylation in the human genome occurs at cytosine-phosphate-guanine (CpG) nucleotides. These have long been considered a repressive mark based on early studies of promoters where methylation correlated with transcriptional repression [1]. Methylation at transcription factor binding sites has previously been thought to correlate with the repression of transcription by either disrupting the binding of methylation-sensitive transcription factors or by having no effect on methylation-insensitive transcription factor binding (Figure 1A) [2]. However, recent high throughput studies have found that methylation within transcription factor binding sites can lead to increased or decreased transcription factor binding dependent on the position within the motif [3,4]. Recent work has shown that the strength of the effect of methylation on transcription factor binding affinity varies between nucleotides within a single transcription factor motif. It is vital to determine the specific functional impact of methylation within transcription factor binding sites as aberrant methylation is a hallmark of many human diseases, including cancer, schizophrenia, and autism spectrum disorders [5–7]. Methods that can better predict these effects on gene transcription can assist in identifying and prioritizing potentially harmful variations.

**Figure 1.**
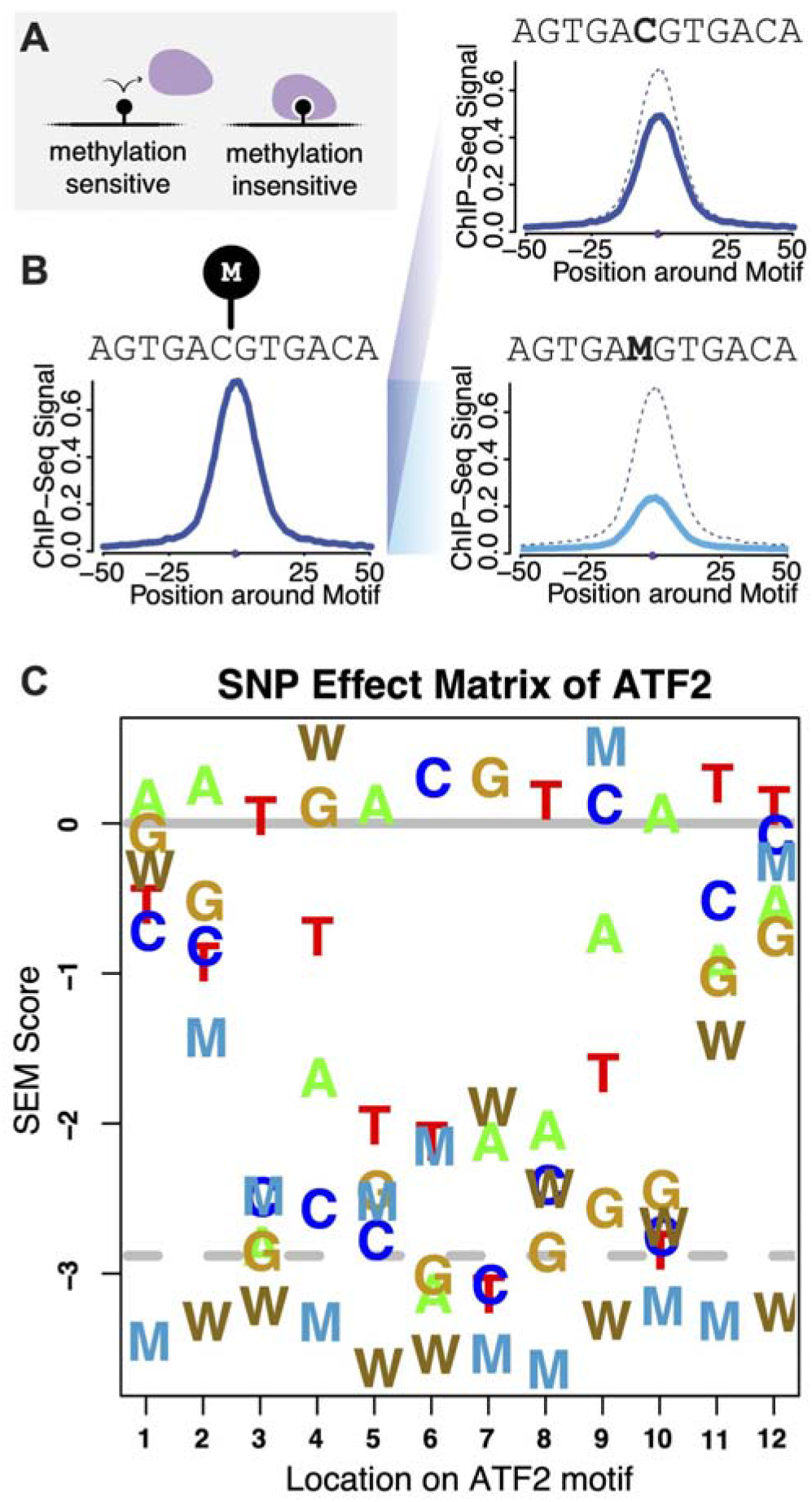
SEM pipeline with methylation predicts the effect of methylation on transcription factor binding affinity. A. Traditional model of methylation sensitivity where methylation sensitive transcription factors are unable to bind their site with DNA methylation present, and methylation insensitive transcription factors can bind regardless of the presence of DNA methylation. B. SEMplMe expands on SEMpl output by adding WGBS to divide ChIP-seq signal peaks of C and W into the proportion of their signal affected by DNA methylation using a weighted sum. C. SEMplMe output is displayed as all 6 nucleotides, including methylated C (M), and G opposite to methylated C (W), at every position along the motif. All values are displayed as log 2 and normalized to an endogenous binding baseline set to 0 (dark gray line). A scrambled baseline is also included (dashed gray line). SEM shown

The effect of methylation on the binding of individual proteins has been studied *in vitro* using protein binding microarrays (PBMs) and newer systematic enrichment of ligands by exponential enrichment (SELEX) based methods [8–11]. Both PBMs and SELEX rely on proteins binding to DNA fragments *in vitro* and may not recapitulate endogenous binding patterns within the genome. A recent study of methylation sensitivity in 542 human transcription factors using a high throughput SELEX method, methyl-SELEX, found 23% of transcription factors were sensitive to methylation, 34% were enhanced by methylation, and 40% were insensitive to methylation [3]. Computational methods to analyze methyl-SELEX data, such as Methyl-Spec-Seq, provide quantitative information on the magnitude and direction of the predicted effect of methylation on transcription factor binding [12]. Additional SELEX-based studies have also observed differences in methylation sensitivity between different positions in a single motif, and is supported by evidence that some bases within transcription factor binding motifs are more correlated with disease compared to others [13,14]. However, predictions produced from these methods are limited as they rely on *in vitro* SELEX data and may not reflect binding patterns in a native context. Methods to determine the methylation sensitivity of transcription factors *in vivo* exist, however they are experimentally rigorous, or do not directly estimate methylation consequence on transcription factor binding, and are therefore challenging to use for broad interpretation [4,15,16]. A robust method to study the native context of DNA methylation within transcription factor binding sites using *in vivo* data is still needed to more accurately model the role these epigenetic marks play on transcriptional regulation.

To address this need, we have adapted SEMpl, a computational genome-wide transcription factor binding affinity prediction method, to incorporate whole genome bisulfite-seq (WGBS) data. This allows our predictions to include the effects of DNA methylation on binding affinity [17]. SEMpl uses open-source *in vivo* data to generate predictions using transcription factor binding data from ChIP-seq and open chromatin data from DNase-seq for a transcription factor of interest. The results are displayed as a SNP effect matrix providing predictions for every potential base change in a transcription factor’s motif. Our SNP Effect Matrix pipeline with Methylation (SEMplMe) method expands these results by incorporating methylation data from WGBS, generating predictions that encompass the magnitude and direction of change to transcription factor binding for all 4 nucleotide base pairs, and adds two additional nucleotide letters: methylated C (M), and G opposite to a methylated C (W). This new tool provides improved specificity to determine which variants lead to disruption of transcription factor binding by integrating endogenous functional information on methylation states and transcription factor binding, advancing our ability to interrogate and prioritize mutations likely to be associated with human disease.

## Methods

### Usage/accessibility

SEMplMe is open source and can be downloaded from https://github.com/Boyle-Lab/SEMplMe. Precomputed SEMplMe plots are available for more than 70 transcription factors (Supplementary Table 1).

### SNP Effect Matrix pipeline with Methylation

SEMplMe functions as an extension of our previously published expectation-maximization method, SEMpl [17]. SEMpl uses an estimation-maximization-like algorithm to predict the consequence variation to binding in transcription factor binding sites using three data types: chromatin immunoprecipitation followed by deep sequencing (ChIP-seq) data, endogenous measures of transcription factor binding genome-wide, DNase I hypersensitive site (DNase-seq) data, a measure of open chromatin genome-wide where transcription factors are known to function; and position weight matrices (PWMs), representing previous knowledge of the binding motif for a transcription factor. SEMpl uses these data to estimate the consequence of all possible variants in a binding site and outputs a matrix of predictions for each of the four nucleotides at each position of a transcription factor’s motif. By including whole genome bisulfite sequencing (WGBS) data to the final output of SEMpl, we have expanded the interpretation of our algorithm to include the contribution of DNA methylation on transcription factor binding (Figure 1B). Starting PWM and cell type used to generate this SEM and all other data shown in figures in this paper can be found in Supplementary Table 2.

In order to evaluate the consequence of DNA methylation in transcription factor binding sites we first gathered WGBS for each kmer aligned to the genome containing an *in silico* SNP. All data shown was generated using matched cell types for ChIP-seq, DNase-seq, and WGBS data. As the vast majority of sites in WGBS data methylation are not binary, the contribution of the proportion of methylation on binding for C and G SNPs at each position within a motif is calculated. Methylation is calculated for each aligned SNP list using the equation: 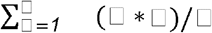, where M represents the proportion of methylation for an aligned kmer, S represents the ChIP-seq signal for it’s alignment, and n represents the total number of kmers in the list. Therefore, the equation: 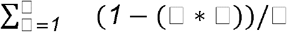 represents the signal contribution of the non-methylated kmer. Using this method, cytosines are divided into methylated and non-methylated components for each position within the motif of a transcription factor. Following this, all 6 nucleotides are included in a SNP effect matrix at each position along the motif of the transcription factor and plotted for an easy to visualize model of transcription factor binding (Figure 1C).

SEMplMe is written in C++, perl and R. In addition to a the matrix file (.me.sem) and the pdf of the visualized sem (*_semplot.me.pdf), the output also includes a matrix of standard error (.sterr) and a matrix of total ChIP-seq signal (.me.totals). New alignment and baseline files are also generated for SEMplMe (.me). A quality control file was used, which provides the −log10(P-value) of the average of 100 *t*-tests from 1000 randomly chosen kmers from the signal files versus the scrambled signal files from SEMpl (Supplementary Table 1). A threshold of 3.15 was set to report confidence in a SEM plot, with runs falling under this threshold highlighted in red.

### SEMplMe sequence scoring

Scoring a full sequence with SEMplMe can be done in the same manner as PWMs or SEMpl, where the log2 score analogous to the nucleotide of interest at each position is added to reflect the predicted binding score of the sequence. This allows predictions to be made for motifs carrying more than one variant.

### EMSA

Kd values for CEBPB and ATF4 were calculated from a previously published EMSA reaction by densiometric scanning by ImageJ and the Excel Solver Package [9,18,19]. All EMSA scores are represented as a ratio to the unmethylated control.

### Correlation with ChIP-seq data

All kmers likely to bind CTCF were recovered from the final iteration of SEMpl. For each kmer with at least 50 occurrences, the average ChIP-seq signal and standard error were calculated. Correlation cutoffs for SEMplMe were defined as the scrambled baseline for the final iteration of SEMpl.

## Results

### SEMplMe provides quantitative predictions based on *in vivo* measures of binding affinity

SEMplMe integrates endogenous functional data encompassing transcription factor binding, open chromatin, and DNA methylation to provide a quantitative prediction of the effect of methylation on transcription factor binding affinity at every position within a binding motif. By including measures of DNA methylation, SEMplMe is able to calculate the relative average transcription factor binding affinity of methylated genomic sequence by using a weighted sum of ChIP-seq signal and the proportion of methylation at the site from WGBS (Figure 1B). Averaging this signal genome-wide for methylated and unmethylated sequence separately allows for the generation of a quantitative prediction matrix of the effect methylation has on transcription factor binding affinity (Figure 1C). SEMplMe represents an advancement over currently existing methods as its predictions are generated from *in vivo* functional data, it is generally accessible without additional experimental work, and the resulting matrix is both quantitative for a single position and across an entire motif.

### SEMplMe recapitulates differences in methylation sensitivity between transcription factors

Transcription factor differences in methylation sensitivity were examined by calculating the absolute difference between methylated and unmethylated bases at each position within SEMplMe matrices for methylation sensitive and insensitive transcription factors. Methylation sensitive transcription factors examined here include CREB, cMYC, USF, NFkB, E2F, MYC, and ZFX [2,11,20,21]. Methylation insensitive transcription factors examined here include SP1, REST, CEBPa, FOXA1, RXRA, and ARNT2 [2,8,11,21–23]. As expected, transcription factors previously associated with methylation sensitivity show a larger average difference in SEM scores between C and M, and G and W nucleotides compared to transcription factors previously defined as insensitive (Figure 2). This suggests that prior definitions of methylation sensitivity and insensitivity may reflect general trends of transcription factor methylation sensitivity. However, it remains unclear if this trend is driven from methylation sensitivity across an entire motif, or typically driven by a single position.

**Figure 2.**
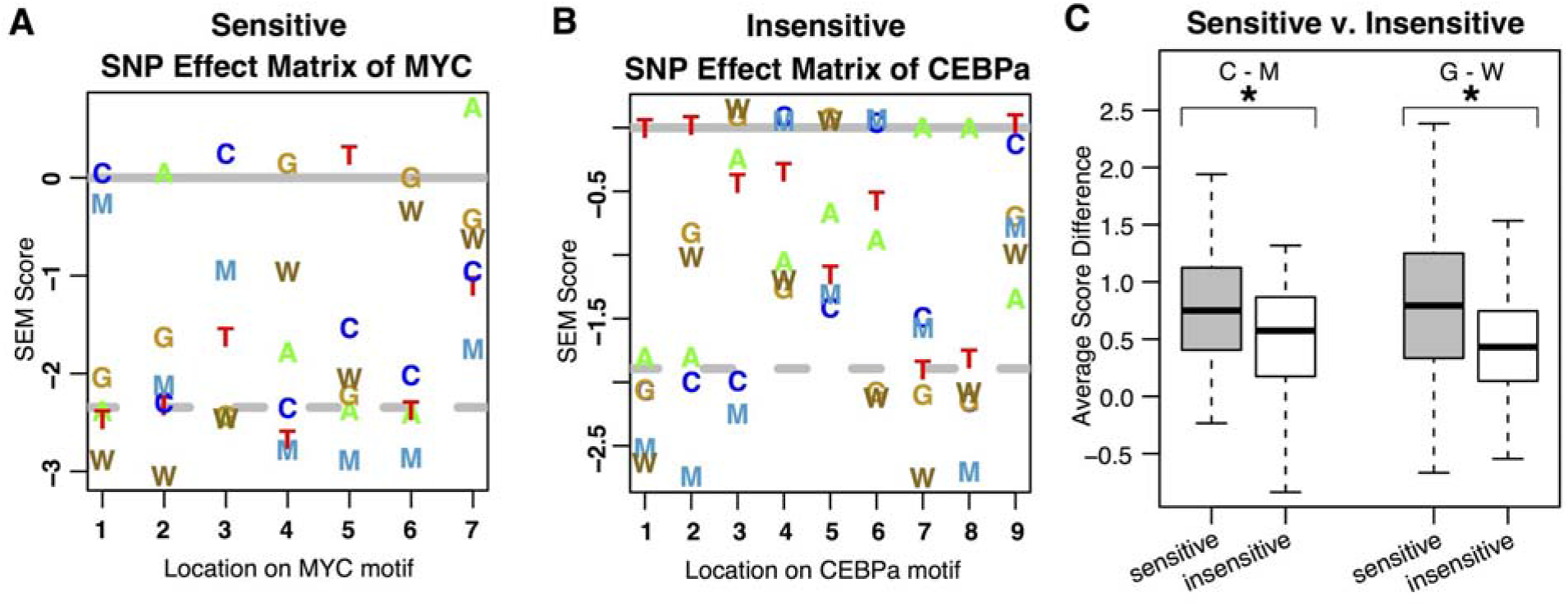
SEMplMe confirms differences in methylated SEM scores for sensitive versus insensitive transcription factors. A. Known methylation sensitive transcription factor MYC shows a large difference between methylated and unmethylated nucleotides at most positions. B. Known methylation insensitive transcription factor CEBPa shows very little difference between methylated and unmethylated nucleotides at most positions. For some positions (i.e. position 5), a small increase in binding is predicted for a methylated cytosine. C. Transcription factors previously annotated as methylation sensitive and insensitive show a significant difference in methylated (M/W) and non-methylated (C/G) SEM scores (T-test C-M P-value = 0.007 and GW P-value = 1.32*10^7). Error bars represent standard deviation.

### DNA methylation drives cell type specific transcription factor binding

DNA methylation is hypothesized to contribute to cell type specific transcription factor binding by altering the availability of DNA sequence. In support of this, the input cell type was found to influence the output of SEMplMe for some transcription factors. One example, JUN, shows high correlation of SEMplMe outputs for methylated sites (MW) between H1-hESC and K562 cell lines (R^2 = 0.91), and a reduced correlation to HepG2 (R^2 => 0.43) (Figure 3). This is supported by MethMotif data, in which JUN shows many more methylated binding sites, most of which fall into a mid-to highly-methylated state in HepG2, as opposed to comparatively few overlapping methylated sites in K562 and H1-hESC [15]. This pattern of reduced correlation was not observed when looking across the entire SEMplMe output, suggesting methylated sites are driving this difference (Supplementary Figure 1). Of note, this pattern is not seen for another transcription factor, CEBPB, where the SEMplMe output for methylated sites is highly correlated between all cell types examined (K562, IMR-90, HepG2, and GM12878), suggesting that not all transcription factors are subject to cell type specificity due to methylation differences (Supplementary Figure 2). Interestingly, SEMpl data without methylation appears to be primarily cell type agnostic, providing evidence that methylation plays a meaningful role in cell type specificity for some transcription factors [17].

**Figure 3.**
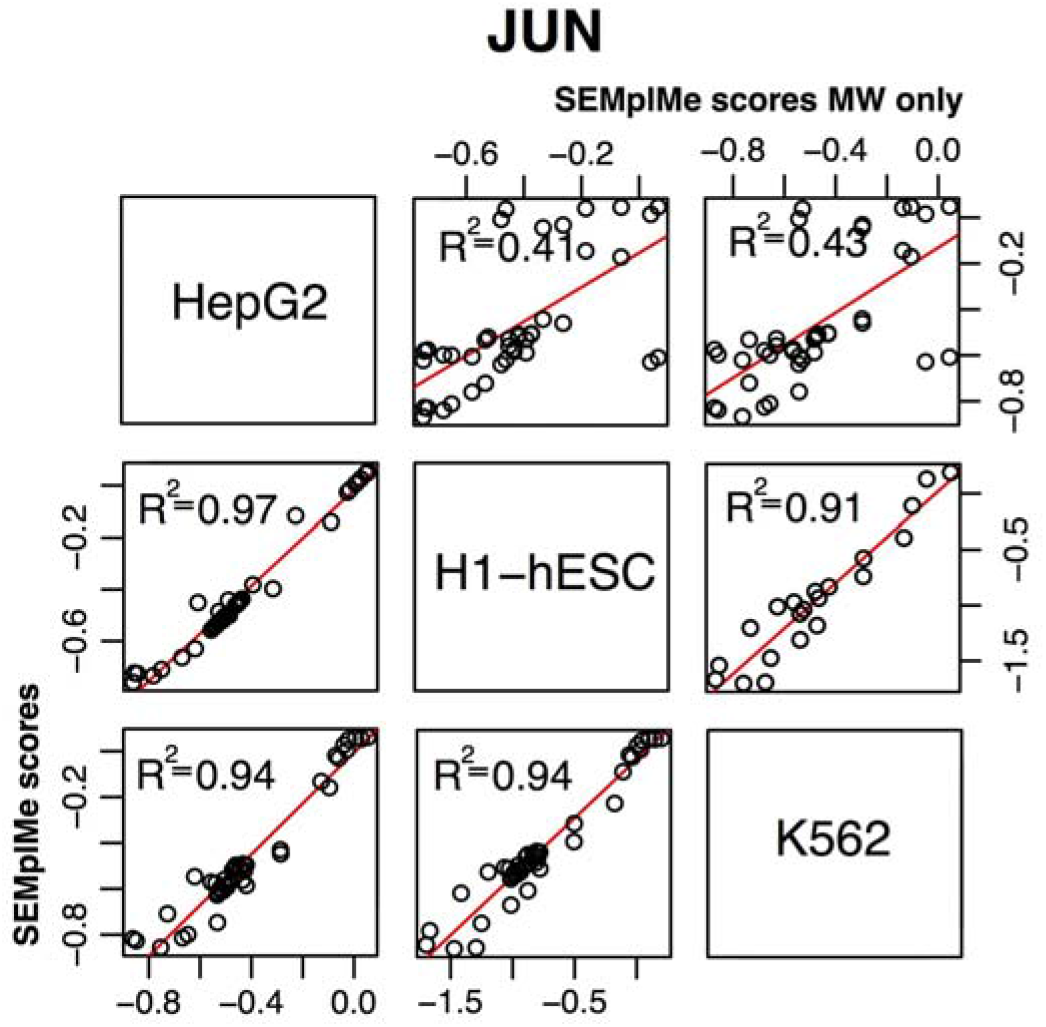
SEMplMe output for JUN varies between cell types. SEM plots vary more between cell types when only considering methylated sites (top right) than methylated and unmethylated sites (bottom left). This suggests methylation plays a key role in the cell type specificity of the transcription factor JUN.

### SEMplMe validation using *in vitro* measures of transcription factor binding affinity

To evaluate SEMplMe on a metric external to ChIP-seq data, our predictions were compared to PBM data, which has been used by previous studies to examine the affinity of individual transcription factors to potential target sequence *in vitro* [8–10]. SEMplMe predictions were compared to microarray Z-scores data from PBMs, which represent transcription factor binding affinity to methylated or unmethylated DNA sequence. A modest level of agreement was observed between SEMplMe predictions and previously published PBM data across 8 transcription factors (Figure 4A)(R^2=0.67)(CEBPA, CEBPB, CEBPD, CREB1, ATF4, JUN, JUND, CEBPG) [10]. This agreement is reduced when using SEMpl scores without methylation (R^2=0.56), suggesting that the inclusion of methylation in our model improves scores for methylated sequences (Figure 4B). Discrepancies between SEM predictions and PBM data can be attributed to differences in *in vivo* versus *in vitro* methods of generation.

**Figure 4.**
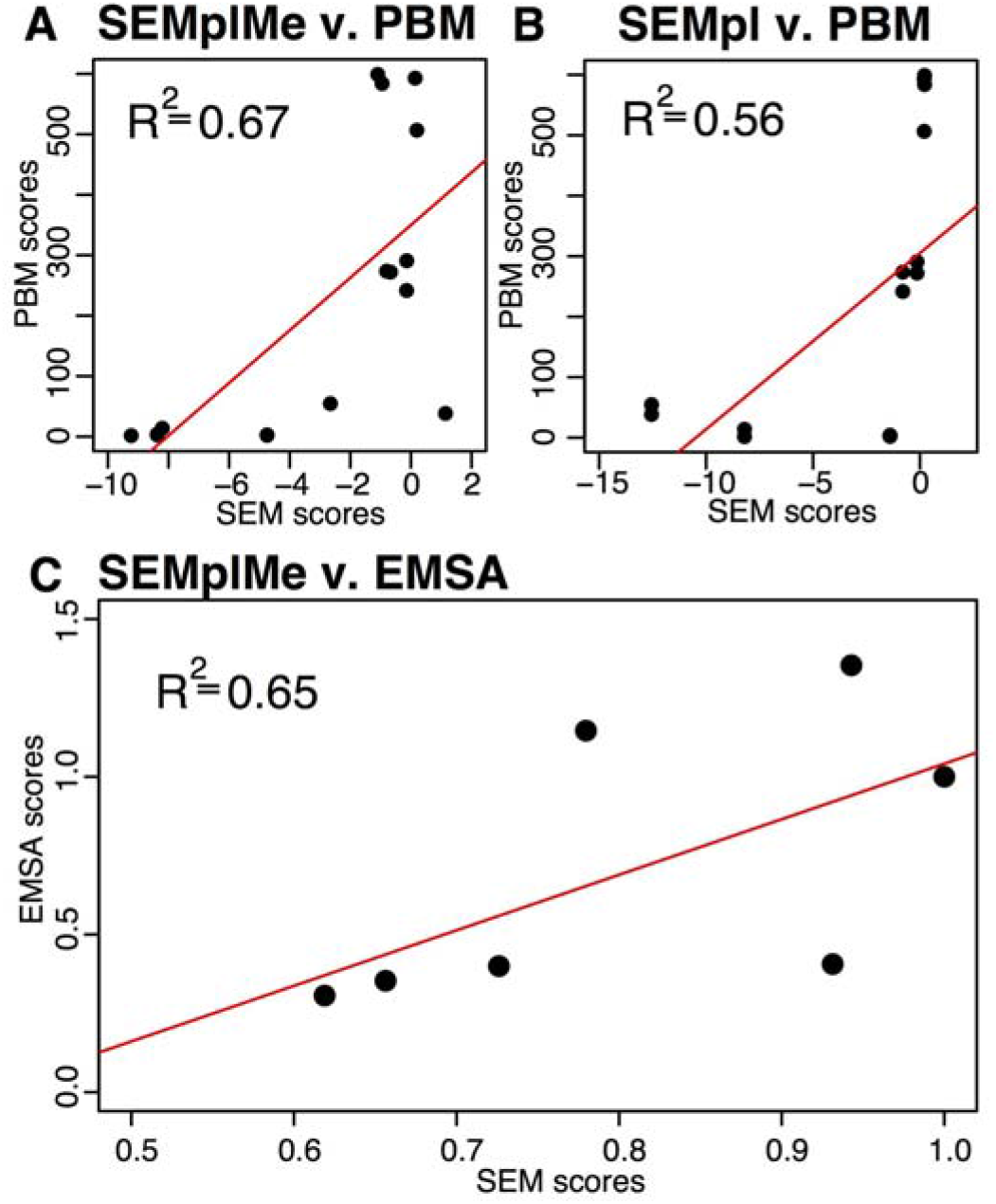
SEMplMe predictions agree with *in vitro* experimental methods. A. SEMplMe agrees with previously published protein binding microarray (PBM) data of methylated and unmethylated binding sites for 8 transcription factors (R^2 = 0.67) [10]. B. SEMpl shows a reduced correlation with PBM data compared to SEMplMe (R^2 = 0.56), suggesting the addition of methylation data improves methylated sequence predictions. C. SEMplMe predictions correlate with previously published electrophoretic mobility shift assay (EMSA) data for methylated, hemi-methylated, and unmethylated binding sites for ATF4 and CEBPB (R^2 = 0.65) [9].

**Figure 5.**
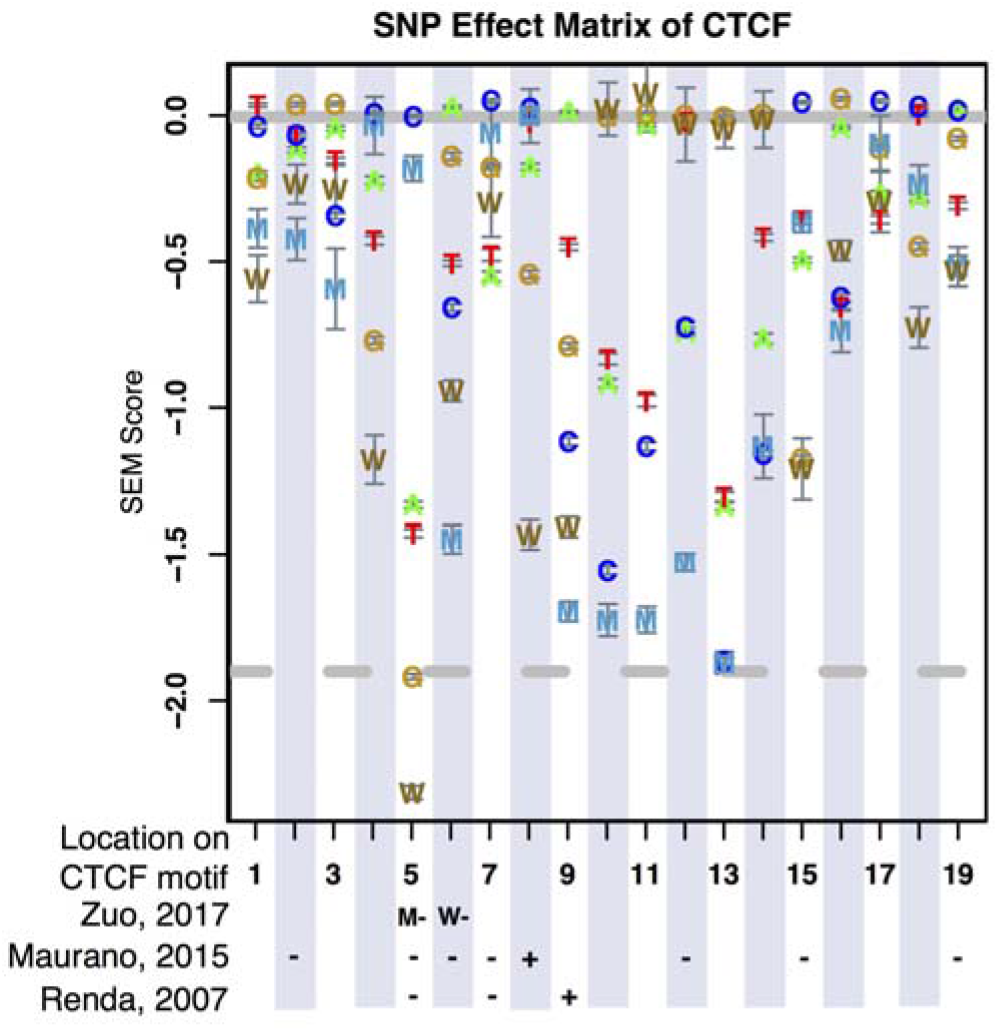
SEMplMe predictions agree with previously published predictions and experimental measures of CTCF binding to methylated sequence. Signs (+/-) found below the SEM plot represent the reduction or increase in binding affinity reported by previous studies at the analogous position. All signs shown without an M or W represent a methylated cytosine (M). Error bars represent standard deviation [4,12,27].

To further functionally validate SEMplMe, data from *in vitro* electrophoretic mobility shift assays (EMSAs) were utilized to examine our predictions. Previously published EMSA data was evaluated for two transcription factors, ATF4(CREB) and CEBPB. This measure of *in vitro* binding showed marginal agreement with our predictions (R^2=0.65)(Figure 4C)[9]. This observed low agreement is driven entirely by CEBPB which has relatively low correlation with our predictions (R^2=0.17), as opposed to ATF4 (R^2=0.92). CEBPB has been reported to preferentially bind to methylated sequence, thus the discrepancy in predictions has previously been thought to be a result of limited genome methylated sequence availability, a necessity for calculating more accurate predictions in SEMplMe [9]. SEMplMe identified comparatively few methylated sites throughout the genome, leading to a much higher standard deviation for the effect of methylated sites (Supplementary Figure 3). This unavailability of methylated sites is consistent with previous data showing methylated CEBPB motifs to bind well *in vitro*, but poorly *in vivo* [24].

### SEMplMe predictions are consistent with previous findings for CTCF

CTCF is a well studied transcription factor previously shown to be methylation sensitive [25,26]. CTCF binding predictions using SEMplMe found the majority of positions to be methylation sensitive for both M and W. Notably, a handful of sites had methylated sequence scores at or slightly above their unmethylated counterpart, and likely represent methylation insensitive positions. These results are consistent with CTCF’s role as a methylation sensitive transcription factor. As CTCF is widely used in research studies, its binding to sites containing methylated positions within its motif have been previously surveyed by a variety of methodologies, including qualitative EMSA, observation of binding following demethylation of cells, and SELEX-based methods [4,12,25,27]. When SEMplMe results were compared to measures of binding at individual positions within the CTCF motif, a general agreement was observed for the direction of binding for all positions predicted to decrease binding affinity (Supplementary Figure 5). Though the majority of sites identified by previous studies within the CTCF motif were found to be overwhelmingly methylation sensitive, two sites were predicted to lead to increased binding affinity when methylated. Though SEMplMe did not identify these positions, one site overlaps a SEMplMe position consistent with methylation insensitivity, and the other was found to not significantly increase binding by all prior studies [4]. Overall, our predictions are consistent with previous studies of CTCF binding direction.

Correlation between the entirety of the CTCF matrices generated by SEMplMe and the recently published Methyl-Spec-seq method, which uses *in vitro* SELEX to predict methylation effects on transcription factor binding affinity, was assayed (R^2=0.56) (Figure 6A)[12]. SEMplMe outperformed Methyl-Spec-seq by performance comparison when comparing scores across entire kmers to their average ChIP-seq signal (SEMplMe R^2=0.25, Methyl-Spec-seq R^2=0.04) (Figure 6B&C). The kmer set used is associated with active CTCF binding and includes both methylated and unmethylated sequences. This provides further evidence that predictions of change to transcription factor binding affinity perform better when generated from *in vivo* data, rather than *in vitro* data such as from SELEX methods.

**Figure 6.**
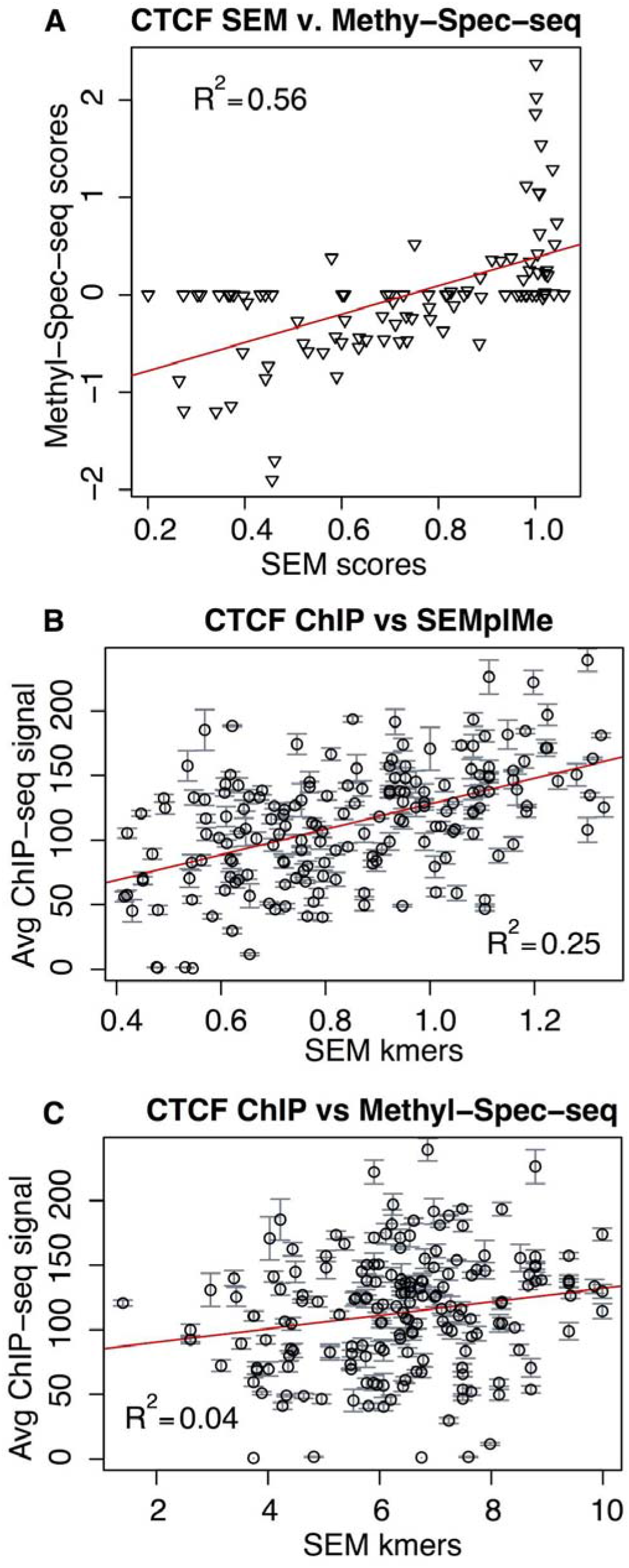
SEMplMe has higher correlation with *in vivo* CTCF binding than Methyl-Spec-seq. A. Correlation of CTCF matrices between SEMplMe and Methyl-Spec-seq show a modest agreement (R^2=0.56). B&C. SEMplMe outperforms Methyl-Spec-seq when comparing to CTCF ChIP-seq scores for whole kmers, including methylated sites (SEMplMe R^2=0.25, Methyl-Spec-seq R^2=0.04).

## Discussion

DNA methylation is a key epigenetic mark known to act in a regulatory capacity, allowing transcription factors to bind in a cell-type specific manner. Counter to the idea that all methylation is able to disrupt transcription factor binding, recent studies have revealed that certain methylated loci impact binding more than others. Predicting the locations of these methylation sensitive loci and quantifying the effect of methylation on transcription factor binding affinity is challenging. Here we introduce an expansion to our previously released software SEMpl, called SEMplMe, which integrates predictions of the effect of cytosine methylation on transcription factor binding affinity based on WGBS data. These predictions agree with *in vitro* data of transcription factor binding, are cell-type specific, and show a general agreement with data from transcription factors previously annotated as methylation sensitive and insensitive.

SEMplMe is poised to advance our understanding of the effects of methylation on transcription factor binding affinity through its generation of quantitative predictions using *in vivo* functional data. SEMplMe will both improve our ability to predict putative disease loci affected by aberrant DNA methylation, and increase predictions of transcription factor binding affinity in general [21]. This is expected to hold true regardless of whether reduced methylation in a transcription factor’s motif contributes to its binding, or is caused by its binding [28]. The nucleotide W was included to capture not just position dependent, but stand dependent methylation, as strand specificity due to hemi-methylation has previously been found to influence transcription factor binding [12]. This is likely driven by changes in DNA structure.

SEMplMe has similar limitations to its predecessor SEMpl, such as a dependence on available ChIP-seq, DNase-seq, and WGBS data. It is further restricted by the limited number of methylated sites in the genome available for use in generating models of binding. In instances where few sequence specific sites also contain methylation, our measure of standard deviation increases considerably. Though the low confidence in these sites can be visualized by error bars, predictions of methylation at these loci are limited. Cell type should be carefully considered before running SEMplMe for optimal predictions as cell type specificity contributes to the final SEMplMe plot, and methylation sensitivity has been previously found to be paralog specific [13]. Additional cell type specific factors such as transcription factor co-binding may also affect the final SEMplMe plot as seen for MYC in SEMpl, though is expected to be rare due to the genome-wide nature of the predictions [17]. Starting PWMs should also be carefully considered for transcription factors known to recognize different methylated and unmethylated motifs [8].

The inclusion of CpG methylation provides additional information to help fully understand context-specific transcription factor binding. However, the addition of more nuanced molecular mechanisms that contribute to transcription factor binding are likely to further improve our predictions. This includes additional types of DNA methylation, such as hydroxymethylation and nonCpG methylation, as well as measures of structural changes to the genome [10,13,29–31].

The improved predictions provided by SEMplMe will contribute to a better understanding of the key positions within transcription factor binding sites affected by DNA methylation. This advancement is central to improving our ability to prioritize mutations associated with aberrant methylation contributing to human disease.

---

**Supplementary Figure 1.**
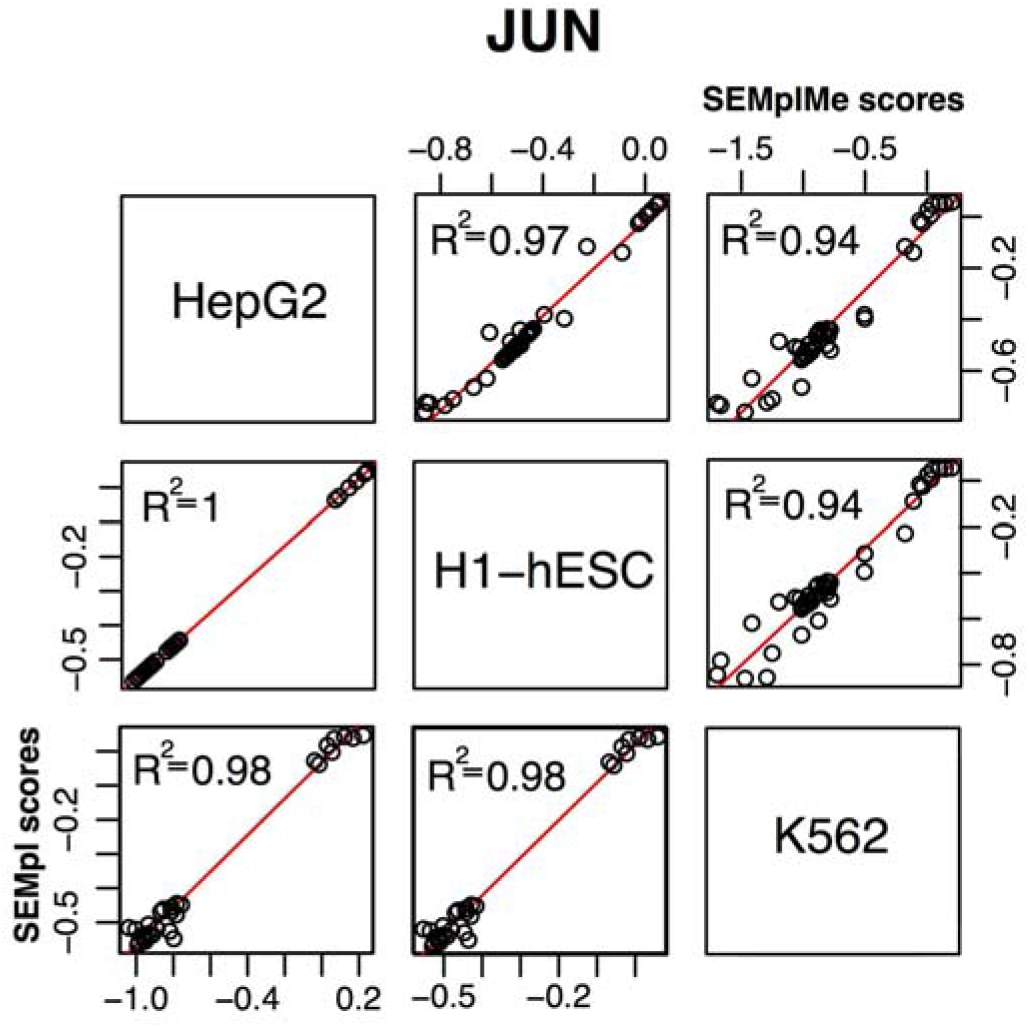
SEMplMe output between cell types versus SEMpl without methylation. SEM plots show more variance between cell types for SEMplMe (top right) than SEMpl without methylation (bottom left).

**Supplementary Figure 2.**
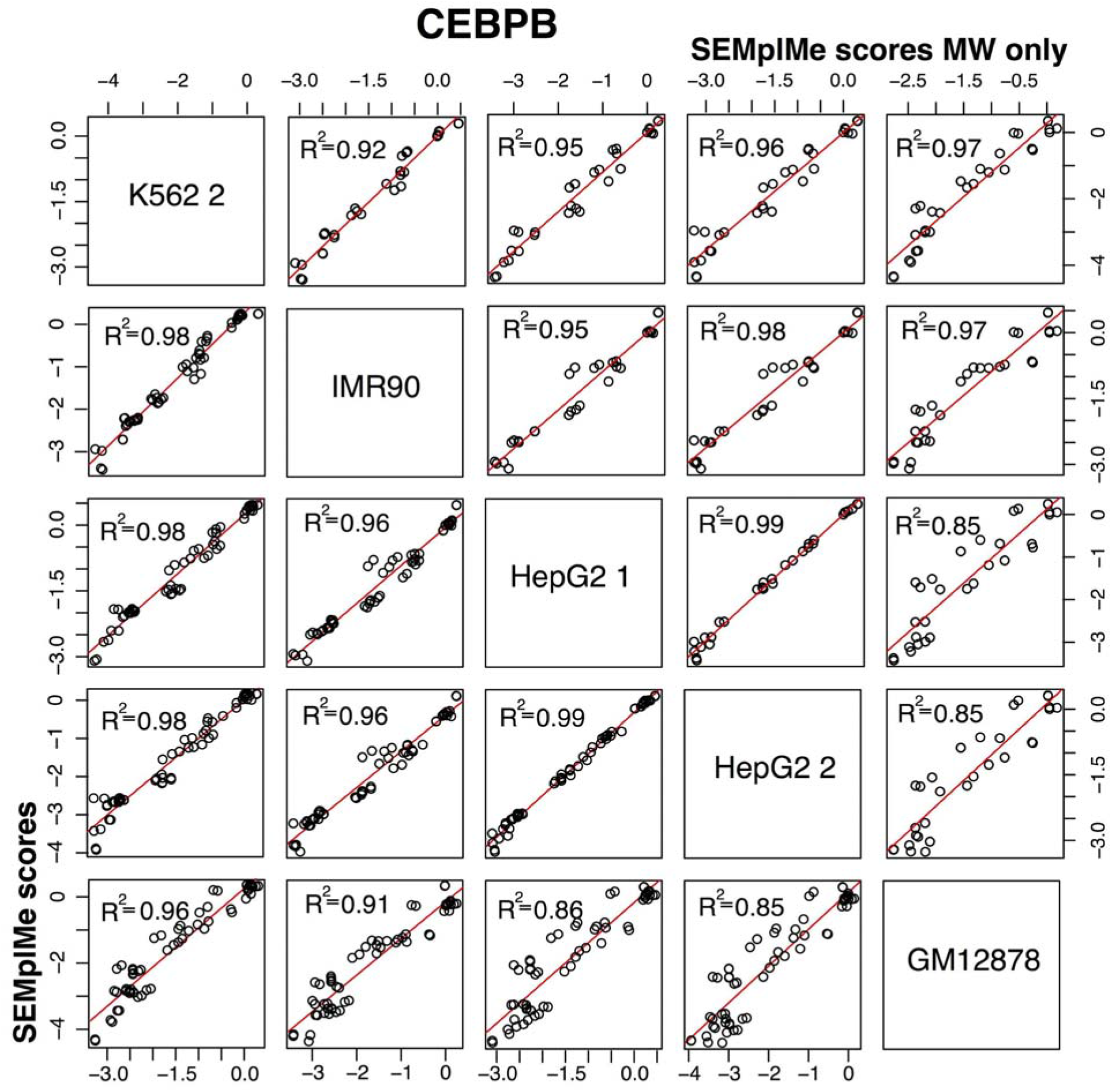
CEBPB SEM output between cell types. SEM plots show little variance between cell types when considering only methylated sites (top right) or both methylated and unmethylated sites (bottom left) for CEBPB. This suggests methylation does not play a large role in cell type specificity for CEBPB.

**Supplementary Figure 3.**
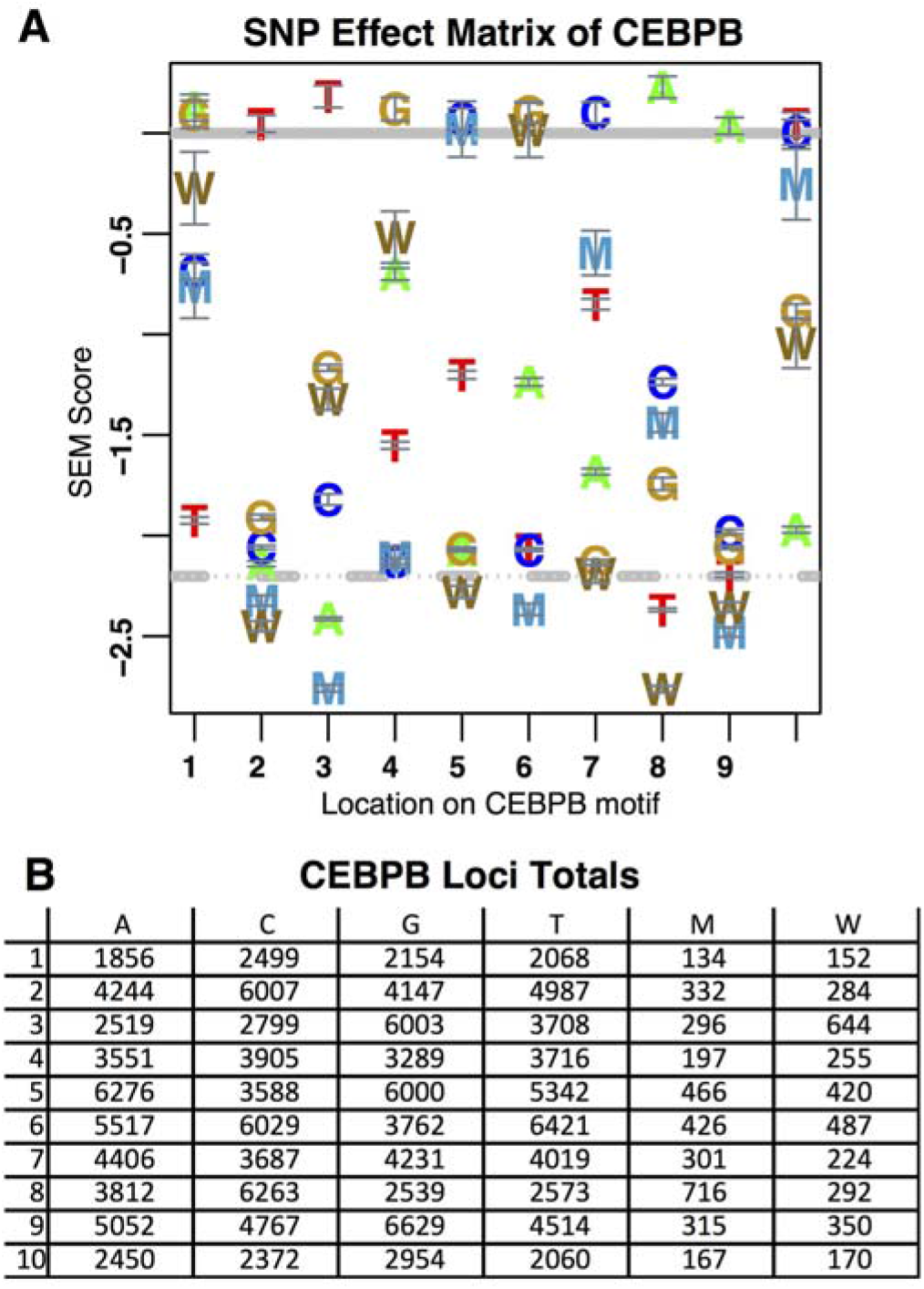
Total number of kmers for each nucleotide in the SEM of CEBPB. A. SEM plot of CEBPB with error bars representing standard deviation. B. Counts of mapped kmers in the genome for each nucleotide at each position. These counts are inversely proportional to the standard deviation seen in the SEM.

