## Supplementary Data for "SEMplMe: A tool for integrating DNA methylation effects in transcription factor binding affinity predictions"

**Supplementary Figure 1.** SEMplMe output between cell types versus SEMpl without methylation. SEM plots show more variance between cell types for SEMplMe (top right) than SEMpl without methylation (bottom left).

**Supplementary Figure 2.** CEBPB SEM output between cell types. SEM plots show little variance between cell types when considering only methylated sites (top right) or both methylated and unmethylated sites (bottom left) for CEBPB. This suggests methylation does not play a large role in cell type specificity for CEBPB.

**Supplementary Figure 3.** Total number of kmers for each nucleotide in the SEM of CEBPB. A. SEM plot of CEBPB with error bars representing standard deviation. B. Counts of mapped kmers in the genome for each nucleotide at each position. These counts are inversely proportional to the standard deviation seen in the SEM.

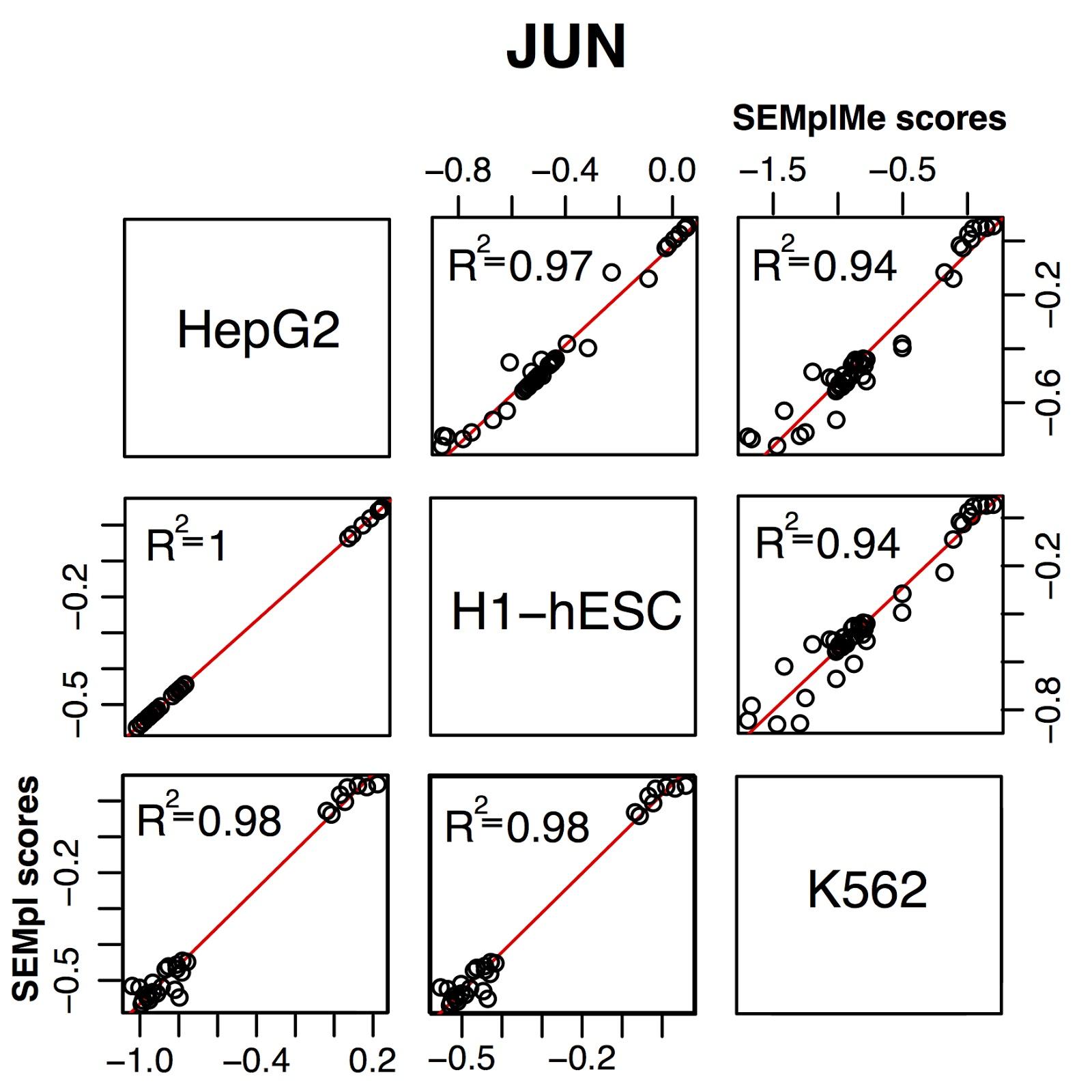

Supplementary Figure 1. SEMplMe output between cell types versus SEMpl without methylation. SEM plots show more variance between cell types for SEMplMe (top right) than SEMpl without methylation (bottom left).

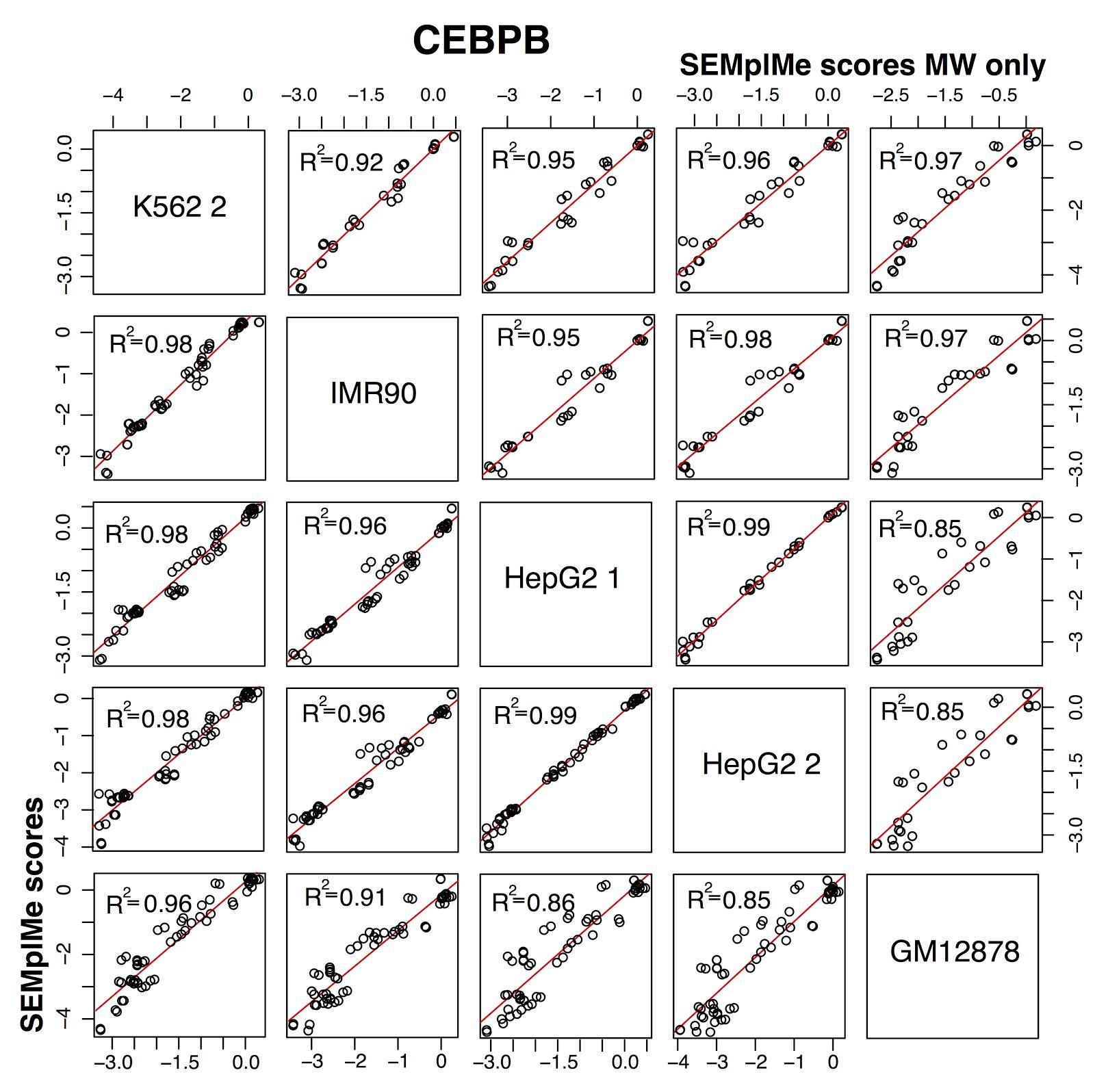

Supplementary Figure 2. CEBPB SEM output between cell types. SEM plots show little variance between cell types when considering only methylated sites (top right) or both methylated and unmethylated sites (bottom left) for CEBPB. This suggests methylation does not play a large role in cell type specificity for CEBPB.

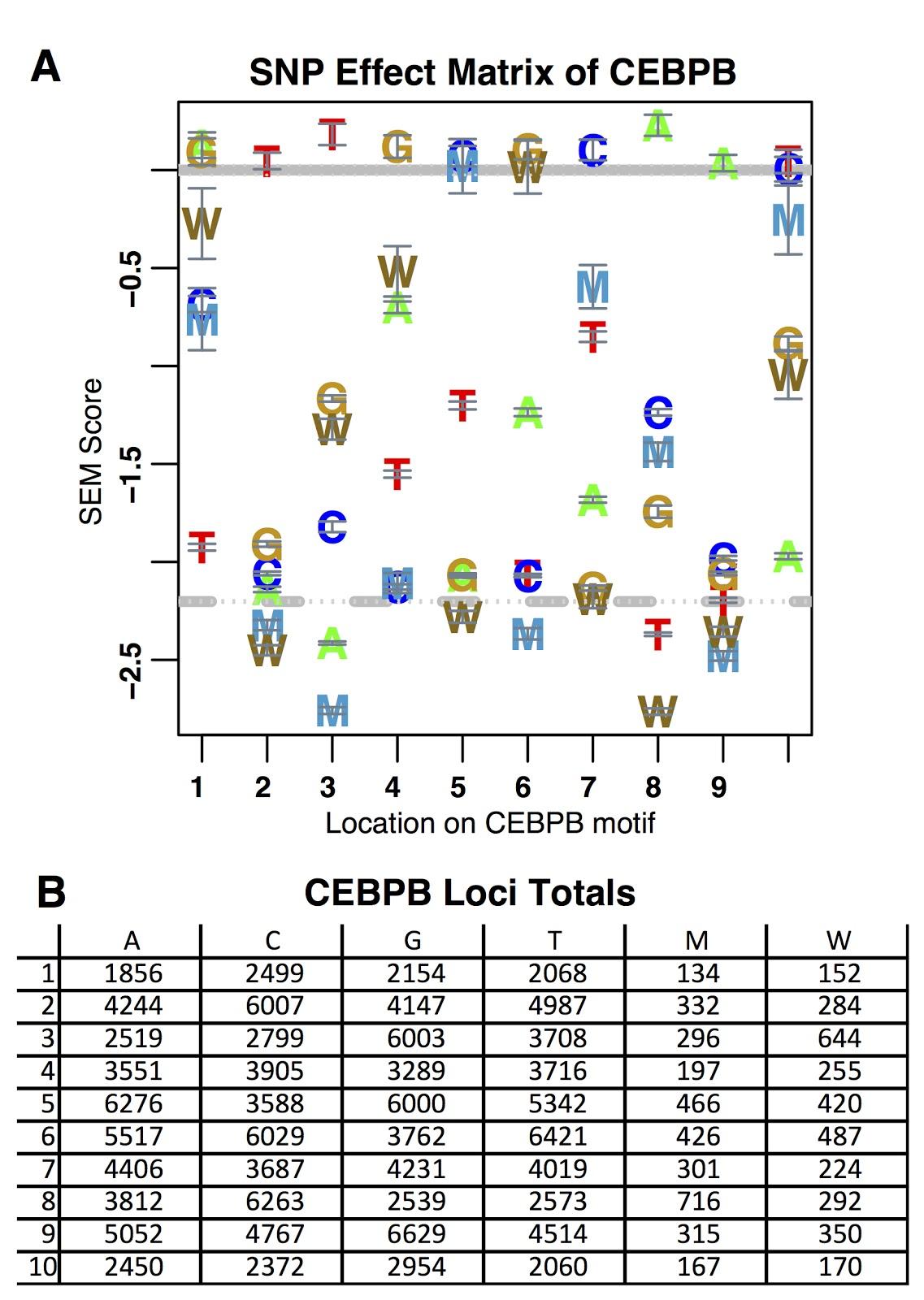

Supplementary Figure 3. Total number of kmers for each nucleotide in the SEM of CEBPB. A. SEM plot of CEBPB with error bars representing standard deviation. B. Counts of mapped kmers in the genome for each nucleotide at each position. These counts are inversely proportional to the standard deviation seen in the SEM.
